# Bipolar oscillations between positive and negative mood states in a computational model of Basal Ganglia

**DOI:** 10.1101/205310

**Authors:** Pragathi Priyadharsini Balasubramani, V. Srinivasa Chakravarthy

**Affiliations:** University of Rochester Medical Center, Department of Neuroscience, Rochester, New York, 14642; Bhupat and Jyoti Mehta school of Biosciences, Department of Biotechnology, Indian Institute of Technology- Madras, Chennai - 36

**Keywords:** cortico-basal ganglia network, serotonin dysfunction, dopamine dysfunction, reward hypersensitivity, basal risk sensitivity

## Abstract

Bipolar disorder is characterized by mood swings - oscillations between manic and depressive states. The swings (oscillations) mark the length of an episode in a patient’s mood cycle (period), and can vary from hours to years. The proposed modeling study uses decision making framework to investigate the role of basal ganglia network in generating bipolar oscillations. In this model, the basal ganglia system performs a two-arm bandit task in which one of the arms leads to a positive outcome, while the other leads to a negative outcome. In healthy conditions, the model chooses positive action and avoids negative one, whereas under bipolar conditions, the model exhibits slow oscillations in its choice of positive or negative outcomes, reminiscent of bipolar oscillations. The model is cast at three levels of abstraction: 1) a two-dimensional dynamical system model, 2) a phenomenological basal ganglia model, 3) a detailed network model of basal ganglia. Phase-plane analyses on the simple reduced dynamical system with two variables reveal the essential parameters that generate pathological ‘bipolar-like’ oscillations. Phenomenological and network models of the basal ganglia extend that logic, and interpret bipolar oscillations in terms of the activity of dopaminergic and serotonergic projections on the cortico-basal ganglia network dynamics. The network’s dysfunction, specifically in terms of reward and risk sensitivity, is shown to be responsible for the pathological bipolar oscillations. The study proposes a computational model that explores the effects of impaired serotonergic neuromodulation on the dynamics of the cortico basal ganglia network, and relates this impairment to abstract mood states (manic and depressive episodes) and oscillations of bipolar disorder.

## Introduction

### Bipolar oscillations as impaired decision dynamics

Optimal decision making consists of the problem of selecting the best choice from a set of potential alternatives. Rewarding or punitive outcomes can shape future decisions. In psychological terms, rewards and punishments may be thought to represent opposite ends on the affective valence scale. There have been efforts to find dissociable brain systems that code for processing rewarding and punitive outcomes (Liu et al., 2011). However, a stringent division of brain systems into reward and punishment systems was found to be inappropriate since neural correlates of reward often overlap with those of punishment as well (Rogers, 2011).

The science of learning about the environment through outcomes (rewards and punishments), and using the results of such learning for decision making, is called reinforcement learning (RL) (Sutton and Barto, 1998b). We focus on a key area of the brain thought to implement decision making as a function of reinforcement—the basal ganglia (BG) (Joel et al., 2002; Schultz, 1998, 2013). Dysfunctional decision making may be associated with pathological mood states - positive in case of mania and negative in case of depression. Therefore, it is believed that positive or negative mood-states and pathological oscillations between them, as found in bipolar oscillations, can be approached through a decision making framework, wherein the agent tries to choose between utilities of mood states with positive and negative outcomes (Alloy et al., 2015; Hilty et al., 2006).

### Basal ganglia and decision making

Basal Ganglia (BG) is a network of subcortical nuclei, known to be involved in a variety of functions including choice selection, timing, working memory, and motor sequencing (Hausdorff et al., 1998; Humphries and Prescott, 2010; McNab and Klingberg, 2008; Mink, 1996; Redgrave et al., 1999; Tanaka et al., 2004; Yahalom et al., 2004). A prominent approach that has been gaining consensus over the past decade, seeks to describe functions of the BG using the theory of RL (Chakravarthy et al., 2010; Joel et al., 2002). RL theory describes how an artificial agent, animal or human subject learns stimulus-response relationships that maximize rewards obtained from the environment. According to this theory, stimulus-response associations with rewarding outcomes are reinforced, while those that result in punishments are attenuated. Experimental studies showing that the activity of mesencephalic dopamine (DA) cells resembles an RL-related quantity called Temporal Difference (TD) error, inspired extensive modeling work seeking to apply concepts from RL to describe BG functions. Thus RL theory is set to account for the diverse and crucial functions of the BG, in terms of the reward-related information carried by mesencephalic DA centers (Houk et al., 1995; Schultz, 2013).

Classically, the functional anatomy of BG, with major nuclei such as striatum, globus pallidus externa (GPe) and interna (GPi), subthalamic nucleus (STN), is thought to consist of two major pathways viz., the direct pathway (DP in which the input port, striatum, is directly connected to the output port, GPi) and the indirect pathway (IP, in which the input port, striatum, is indirectly connected through STN-GPe to the output port, GPi). Another pathway, the hyperdirect pathway (HDP), connecting the cortex to STN of the BG has also been gaining prominence recently (Albin et al., 1989; DeLong, 1990; Nambu et al., 2002). The functional opponency between the DP and IP is the basis of a number of computational models of the BG, which describe the DP and IP pathways as Go and NoGo respectively, in view of their facilitatory and inhibitory actions on movement respectively (Frank et al., 2004; Redgrave et al., 1999). But the expansion of the Go-NoGo picture to Go-Explore-NoGo picture, that includes the IP as a substrate for exploration, accommodated a much wider range of BG functions in a computational RL framework (Chakravarthy and Balasubramani, 2014; Chakravarthy et al., 2010; Kalva et al., 2012). We also found that the BG models can explain a variety of human behavioral data when choice selection is based on optimizing a utility function encompassing reward mean and variance, as opposed to the more traditional value function alone (Balasubramani et al., 2014, 2015a).

### The current study

In the proposed model, we study the utility function approach in more detail, focusing on the influence of subjective reward and risk sensitivity in the overall choice selection dynamics between positive and negative mood states. It uses a simple task framework, a two-arm bandit problem, consisting of probabilistic positive and negative rewards as outcomes of each of the arms (states); they correspond to the model’s positive and negative mood states, respectively. In the first model (A), we build value, risk and utility functions from classic RL strategies for positive and negative states, and use softmax policy (Sutton and Barto, 1998a) to choose between actions. Then, in model (B), we extend the concepts to a more detailed network model of BG, with abstract activities of D1 receptor-expressing Medium Spiny Neurons (D1 MSN) of striatum for computing value, and that of co-expressing D1 and D2 receptor MSNs for computing risk function. The direct and indirect pathways of BG, encompassing STN, GPe, GPi and thalamus, implement the selection strategy to choose between utilities of positive and negative states. So, both the models (A and B) essentially differ in terms of the ‘actor’, that controls the policy (choice strategy) dynamics, and the ‘critic’ that compute utility. In both the models, we associate putative mechanisms driving the selection dynamics with bipolar-like oscillations between mood states. The bipolar oscillations, it must be noted, are obviously different from the pathological oscillations of STN-GPe dynamics that are exhibited in diseases such as Parkinson’s (Gillies et al., 2002; Weinberger et al., 2009; Willshaw and Li, 2002). While the STN-GPe oscillations are in the range of Hertz, the bipolar oscillations span over months and years. Finally, a reduced dynamical system model (C) consisting of a simple two-variable system, that captures the essential dynamics of both the above models, is presented and the correspondences between the key parameters of different models are discussed.

## Model

The task used in this study is a simple two arm bandit problem; one arm (state) is associated with rewarding outcome, reward (r) = 1, and another state with punitive outcome, r = −1. The outcomes are probabilistic with probability = 0.5. We chose this task as it is simple enough for observing the oscillatory effect in decision making framework between positive and negative states.

### A) Phenomenological model (using Softmax policy)

Here, the choice selection dynamics (actor) is carried out by softmax equation, and the critic component computes the utility associated with positive and negative state outcomes (environment). There are auxiliary, slow dynamics that govern the variation of some of the parameters that are involved in the selection dynamics, as elaborated in the model description below (also refer to figure 1).

### B) BG network model

In this version, a more realistic network model of the BG replaces the softmax rule used in the previous model for making a choice (actor). The network model consists of important BG nuclei such as striatum, STN, GPe and interna GPi and thalamus. Utility associated with network inputs is computed as a function of the responses of the striatal neurons. In this case too, there are auxiliary, slow dynamics that govern the variation of some of the parameters that are involved in the selection dynamics of the network model, as elaborated in the model description below (also refer to figure 1).

### C) Reduced dynamical system model

Lastly, a reduced dynamical system of the choice selection and auxiliary dynamics is presented so as to perform phase-plane analysis, and better understand the conditions under which oscillations between positive and negative states are observed (also refer to figure 1).

**Figure 1:**
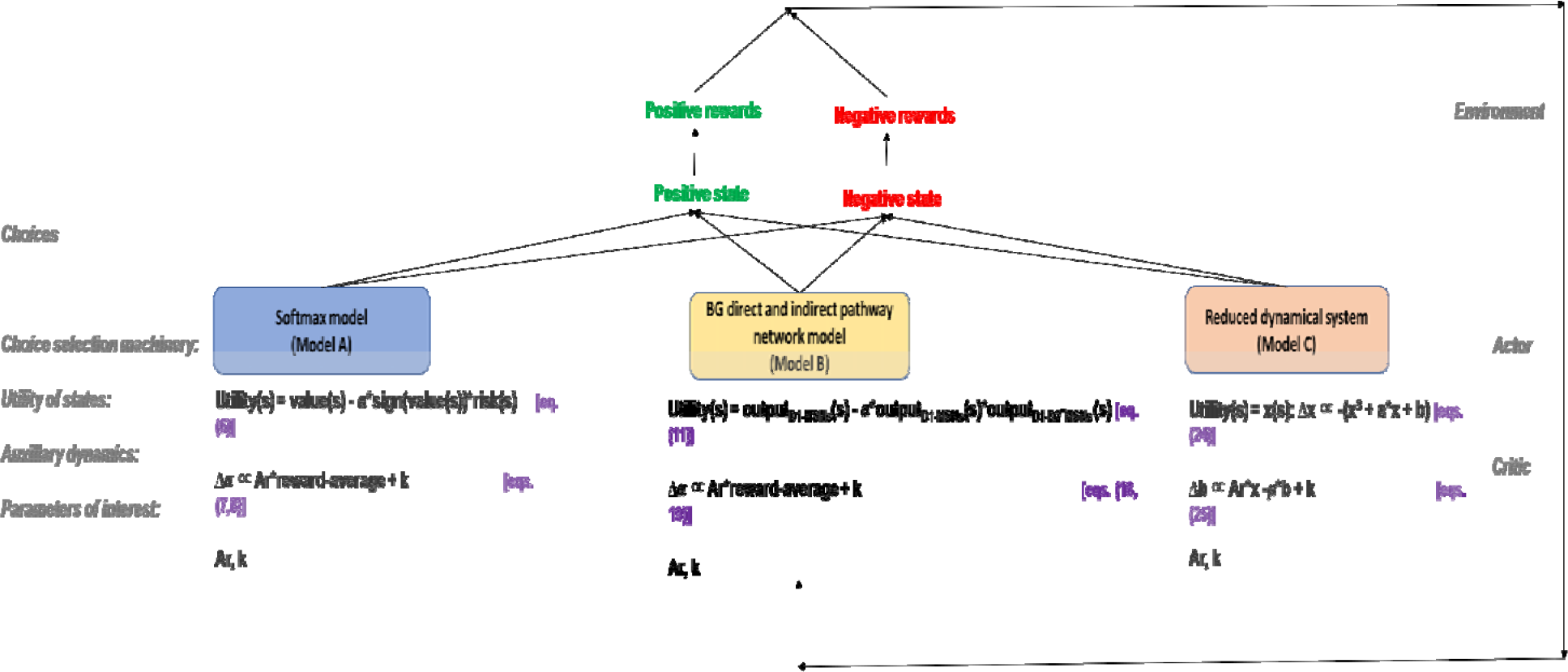
Schematic showing important components of various models used in the study

### Phenomenological model using softmax policy (Model A)

We first present the mathematical formulation for the utility computation in the BG (‘the “critic” component). On the lines of the utility models described in (Bell, 1995) and (d’Acremont *et al.*, 2009), the proposed model of the utility function ‘*U_t_*’ is presented as a tradeoff between the expected payoff and the variance of the payoff (the subscript ′*t*′ refers to time) associated with each state, *s*. The original Utility formulation used in (Bell, 1995; d’Acremont *et al.*, 2009) is given by eqn. (1).

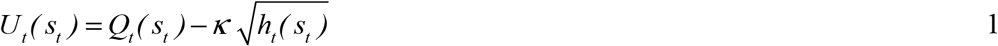

where *Q_t_* is the expected cumulative reward and *h_t_* is the risk function or reward variance, for state, ′*s*′; and ′*κ*′ is the risk preference. Note that in eqn. (1), we represent the state and action explicitly as opposed to that presented in (Bell, 1995; d’Acremont *et al.*, 2009). Following action execution policy ′*π*′, the choice value function ′*Q*′ at time ′*t*′ of a state, ′*s*′, may be expressed as,

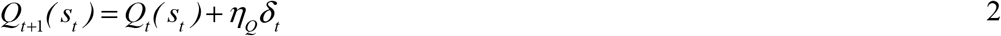

where ′*s*_t_′ is the state at time ′*t*′, ′*a*_t_′ is the action performed at time ′*t*′, and ′*η*_Q_′ is the learning rate of the value function (0 < *η*_Q_ < 1). The temporal difference (TD) error measure of dopamine (DA) is defined by (*δ*_t_ (eqn. 3) for the case of immediate reward problems.

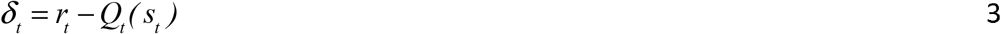

*r*_t_ is the current reward obtained for making a response of choosing state, *s*. Similar to the value function, the risk function ′*h*_t_′ has an incremental update as defined by eqn. (4). Optimizing risk function in addition to the value function (Balasubramani et al., 2014, 2015a) is shown to capture human behavior well in a variety of cognitive tasks involving reward-punishment sensitivity, risk sensitivity, and time scale of reward prediction. The risk function is updated as follows.

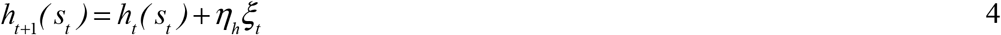

where ′*η*_h_′ is the learning rate of the risk function (0 <*η*_h_< 1), and ′*ξ*_t_′ is the risk prediction error expressed by eqn. (5),

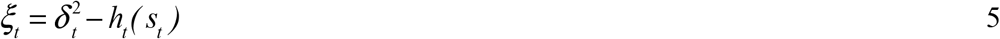

The parameters *η*_h_ and *η*_Q_ are set to 0.01, and *Q*_t_ and *h*_t_ are set to zero at t = 0 for simulations described in the results section. We now present a modified form of the utility function by substituting *κ* = *α sign*(*Q_t_*(*s_t_*, *α_t_*)) in eqn. (1), whose reasoning is given below.

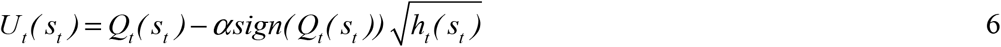

In the above equation, the risk component includes three subcomponents - the ‘*α*’ term, the ′*sign*(*Q_t_*)′ term, and the risk term ′√*h_t_*′. The *sign(Q_t_)* term achieves a familiar feature of human decision making viz., risk-aversion for gains and risk-seeking for losses (Kahneman, 1979; Markowitz, 1952). In other words, when *signa(Q_t_)* is positive (negative), *U_t_* is maximized (minimized) by minimizing (maximizing) risk. Note that the expected choice value *Q_t_* would be positive for gains that earn rewards greater than a reward base (= 0, in the present case), and would be negative otherwise during losses. Our earlier studies have proposed the parameter *α* to be a substrate for serotonin (5-HT) activity in BG (Balasubramani et al., 2014, 2015a).

Updating *α* as a function of rewards is governed by the following equations. This is proposed to represent the adaptation of risk preferences as a function of overall reward averages (Kranz et al., 2010), and they represent the main auxiliary dynamics driving the computation of utility.

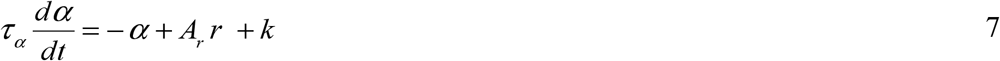

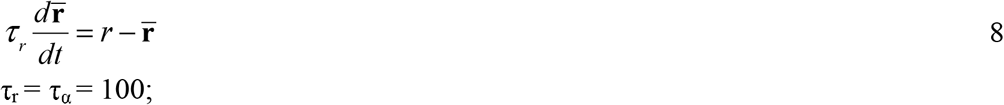

The variable r◻ tracks the average rewards ‘*r*’ gained through time, and the change in *α*, *dα*/ *dt* characterizes the 5-HT dynamics (eqns. 7, 8). The parameter *A*_r_ denotes the reward sensitivity, and thus the reward history modulates *α* dynamics. Eqn. (7) simply means that higher average reward in the recent past must tend to increase risk sensitivity, *α*. Note that the dynamics of *α* is thought to occur at a much longer time scale than that of action selection. ‘*k*’ is a constant.

Choice selection (the “actor” component) is performed using softmax distribution (Sutton, 1998) generated from the utility. Note that traditionally the distribution generated from the choice value is used. The probability, *P_t_(α|s)*, of selecting a choice of state ′*s*′, at time ′*t*′ is given by the softmax policy (eqn. (9)).

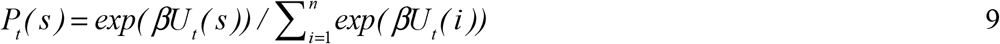

′*n*′ is the total number of choices available, and ′*β*′ is the inverse temperature parameter. Values of *β* tending to 0 make the choices almost equiprobable and the *β* tending to ∞ makes the softmax choice selection identical to greedy choice selection.

### Network model (Model B)

This lumped model of the previous subsection has been extended to the BG network model with the value and the risk functions computed by the medium spiny neurons (MSNs) in the striatum (Balasubramani et al., 2015a; Balasubramani et al., 2015b). Our earlier studies proposed that striatal DA1 receptor (D1R) expressing MSNs code for value function, while the MSNs coexpressing D1R-D2R code for the risk function (Balasubramani et al., 2015b). Whereas the D1R MSNs project via the direct pathway (DP) to GPi, the D2R and the D1R-D2R co-expressing MSNs project to the GPe in the indirect pathway (IP) (Albin et al., 1989; Hasbi et al., 2011; Perreault et al., 2011).

The outputs of the different kinds of MSNs—D1R expressing, D2R expressing and the D1R-D2R co-expressing neurons - are represented by variables *y*_D1_, *y*_D2_, and *y*_D1D2_, respectively in eqn. (10). The subscript *t* denotes the time of response for a particular state, *s*.

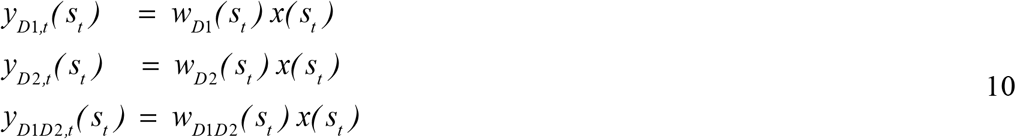

In the above equations, ‘*x*’ is a logical variable modeled to be equal to 1 for the current state, *s*_t_, i.e., *x*(*s*_i_) = 1 *if s*_i_ = *s*_t_. The Utility, *U*, is then obtained from the network model (the ‘critic’ component) as described in eqn. (11) below.

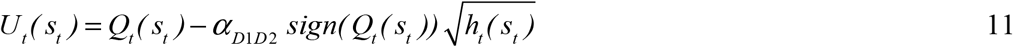

Where

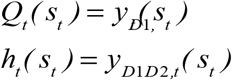

Here in eqn. (11), the risk sensitivity parameter is defined by *α*_D1D2_ that denotes the specific control of 5-HT on the risk function encoding D1R-D2R co-expressing MSNs. In the model, the DA parameter (as described later below) is used for the updating of cortico-striatal weights, and also (as described later below) for controlling the switching at GPi (Chakravarthy and Balasubramani, 2013). The bi-directional connectivity in the STN-GPe system produces complex oscillations and facilitates "exploratory" behavior (Kalva et al., 2012). However, we note that STN-GPe oscillations are different from bipolar oscillations. Whereas the STN-GPe oscillations are in the range of tens of cycles per second, bipolar oscillatory cycles stretch over weeks to years (Alloy et al., 2015; Hilty et al., 2006; Suppes et al., 2000). We now present equations for the individual modules of the proposed network model of the BG contributing to the “actor” component (figure 2). The reader may refer our earlier studies for more modeling details (Balasubramani et al., 2015a; Chakravarthy and Balasubramani, 2014).

**Figure 2:**
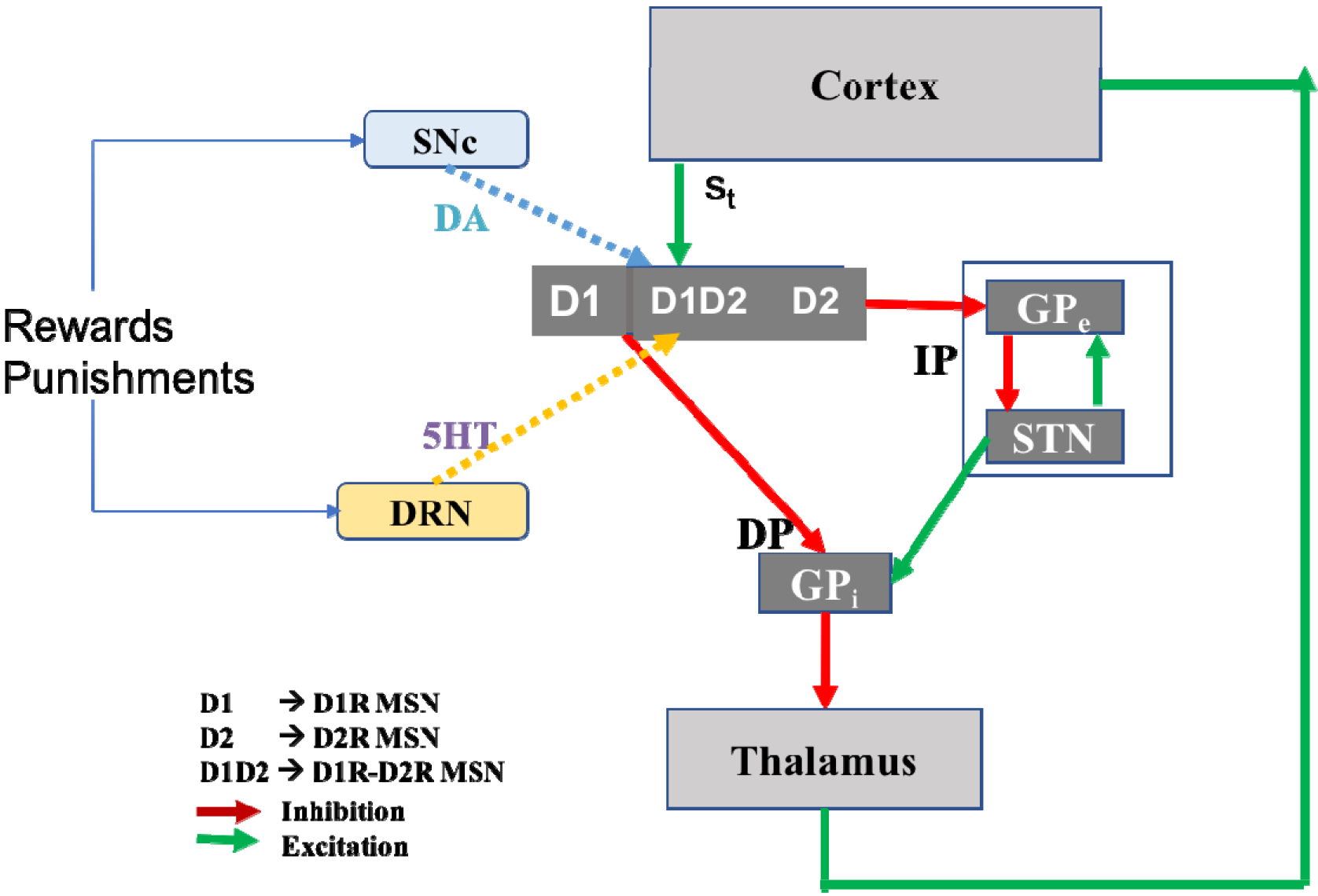
Schematic of the network model of the basal ganglia.

#### Model Components: Striatum

The Striatum is proposed to have three types of MSNs, D1R expressing MSNs, D2R expressing MSNs, and D1R-D2R co-expressing MSNs, all of which have their respective gain functions (*λ*) as described below in eqn. (12). The *c*_1_, *c*_2_, *c*_3_ are constants that vary with the receptor type. The value function (*Q*) of the “critic” module requires a continuously increasing gain as a function of DA in the MSNs, which is shown to occur in the DA D1R containing MSNs. The risk function (*h*) of the “critic” module (Balasubramani et al., 2014, 2015a; d’Acremont et al., 2009)would simply require an increasing gain with increasing *magnitude* of DA, i.e. a ‘U’ shaped gain function which gives increased response with increasing *δ^2^*. It is plausible that these risk-type of gain functions would then probably be exhibited by the neurons that co-express both the D1R-like gain function that increases as a function of DA, and D2R-like gain function that decreases as a function of DA (Humphries et al., 2009; Moyer et al., 2007; Servan-Schreiber et al., 1990; Thurley et al., 2008), as identified in a recent experimental study (Allen et al., 2011). The D2R MSN’s gain function whose activity decreases as a function of DA makes them suitable for punishment computation, in opposition to that of the D1R MSNs responding positively to the reward prediction error (DA).

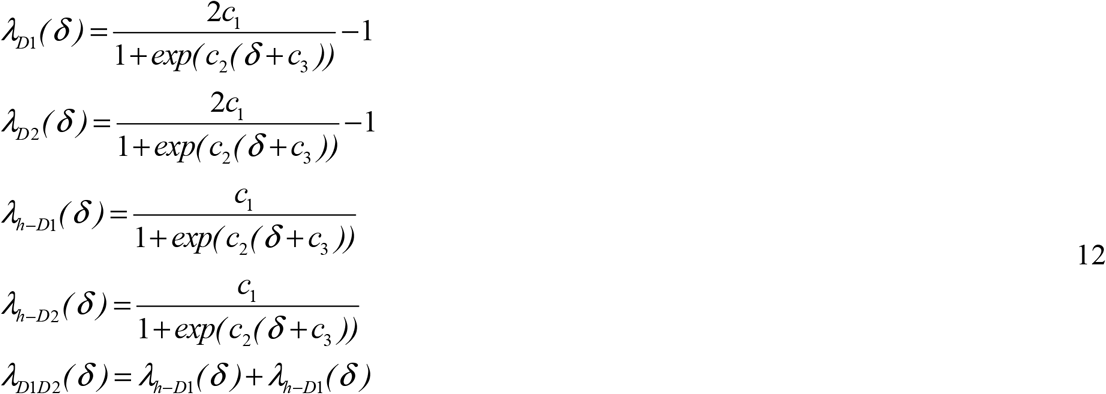

The weight update equations for a given state in the different kinds of MSNs are provided in eqn. (13).

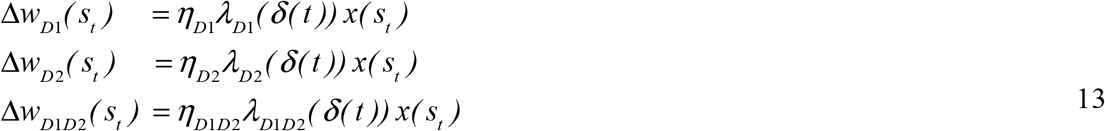

The *δ*’s in the weight update equations are computed for the immediate reward condition as provided in eqn. (14). It represents the DA form of activity that updates the cortico-striatal weights and is the classical temporal difference (TD) error (Houk et al., 2007; Schultz et al., 1997).

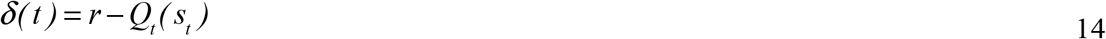

#### STN-GPe system

In the network model of the STN-GPe system, STN and GPe layers have equal number of neurons, with each neuron in STN uniquely connected bidirectionally to a neuron in GPe. Both STN and GPe layers are assumed to have weak lateral connections within the layer. The number of neurons in the STN (or GPe) is taken to be equal to the number of possible choices, viz., positive and negative states, *n* = 2, in our study (Amemori et al., 2011; Sarvestani et al., 2011). The dynamics of the STN-GPe network is given below.

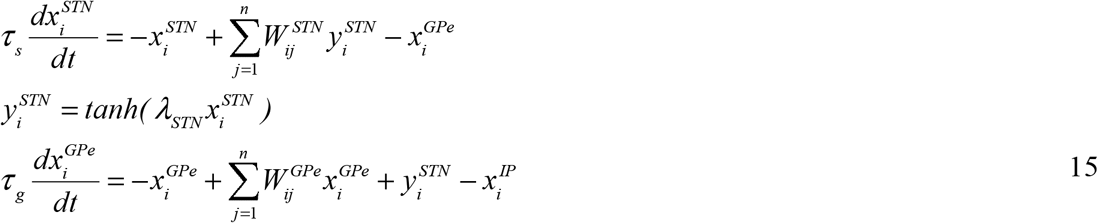

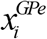: internal state (same as the output) representation of *i*th neuron in GPe;
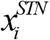: internal state representation of *i*th neuron in STN, with the output represented by 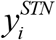;
*W^GPe^*: lateral connections within GPe, equated to a small negative number ◻_g_ for both the self (*i* = *j*) non-self (*i* ≠ *j*) connections for every GPe neuron *i*.
*W^STN^*: lateral connections within STN, equated to a small positive number ◻_s_ for all nonself (*i* ≠ *j*) lateral connections, while the weight of self-connection (*i* = *j*) is equal to 1 + ◻_s_, for each STN neuron *i*.

Both STN and GPe are modeled to have complete internal connectivity, with every neuron in a layer connected to every other neuron with the same connection strength. That common lateral connection strength is ◻_s_ for STN, and ◻_g_ for GPe. Likewise, STN and GPe neurons are connected in a one-to-one fashion – *i*^th^ neuron in STN is connected to *i*^th^ neuron in GPe and vice-versa. For all the simulations presented below, we set ◻_g_ = -◻_s_; the time constants *τ*_S_ = 10; *τ*_g_ = 30.33; and the slope λ_STN_ = 3; ◻_s_ = 0.12.

#### The DP and IP projections to GPi

The outputs of D1R expressing MSNs, transmitted over the direct pathway are computed as:

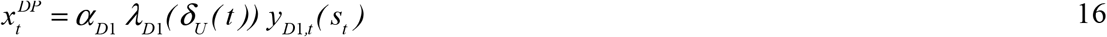

The outputs of the D2R and D1R-D2R expressing MSNs, transmitted to GPe via the indirect pathway, are computed as,

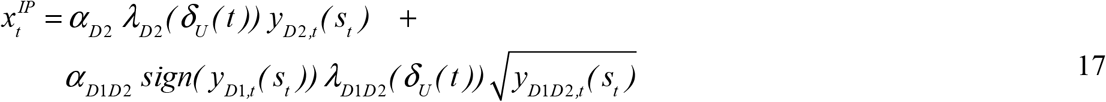

The variables *y*_D1,t_, *y*_D2,t_,*y*_D1D2,t_ as a function of state, *s* at time, *t*, are obtained from eqn. (10). The neuromodulator 5-HT’s specificity in expression along with a particular type of MSN is not known (Eberle◻Wang et al., 1997; Nadjar et al., 2006; Surmeier et al., 1996; Ward and Dorsa, 1996). In the present model, 5-HT is thought to modulate the activity of all three kinds of MSNs (D1R expressing, D2R expressing and the D1R-D2R co-expressing). Hence the modeling correlates of 5-HT are the parameters α_D1_ (eqn. (16)), α_D2_, α_D1D2_ (eqn. (17)) for modulating the output of the D1R, D2R and the D1R-D2R MSNs respectively, and they may represent the 5-HT control exerted by dorsal raphe nucleus (DRN) (Alex and Pehek, 2007; Jiang et al., 1990; Nakamura, 2013). This study allows all 5-HT-related parameters (α_D1_, α_D2_, α_D1D2_) to take the same value, for simplicity (α = α_D1_ = α_D2_ = α_D1D2_). Furthermore, we incorporate 5-HT dynamics as a function of mean observed rewards through time as follows, as there has been considerable evidence suggesting the modulation of 5-HT signaling as a function of rewards (Kranz et al., 2010).

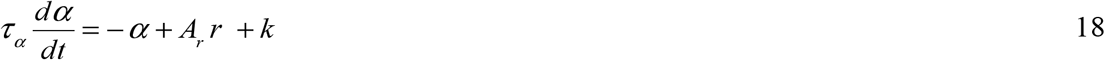

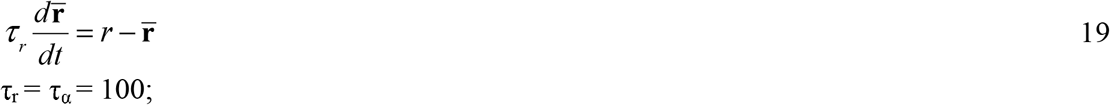

In the above equations, the variable r◻ tracks the average reward gained through time, change in α, *dα/ dt*, characterizes the 5-HT dynamics as in eqns. (7, 8). The parameter *A*_r_ denotes the reward sensitivity. ‘*K*’ is a constant input, is proposed to denote the tonic levels of 5-HT. The last two equations (18, 19) distinguish the network model of (Balasubramani et al., 2014, 2015a) from the network model described in this section, and they form the auxiliary dynamics controlling the model.

The D2R and the D1R-D2R MSNs form part of the striatal matrisomes known to project to the indirect pathway, while the D1R MSNs project to the direct pathway (Amemori et al., 2011; Calabresi et al., 2014; Jakab et al., 1996; Nadjar et al., 2006; Surmeier et al., 1996). It should also be noted that λs used as a gain factor in eqns. (16, 17) have different values from λs used in eqn. (13). The gain functions in eqns. (16, 17) are a function of the DA form (Stauffer et al., 2014) which represents the temporal difference in utility function, *δ*_U_(eqn. 20). This is different from the DA form, *δ*, described in eqn. (14).

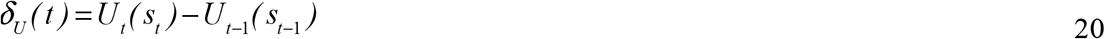

#### Choice Selection at GPi

Choice selection at GPi is implemented using the combination of the DP and IP contributions as follows:

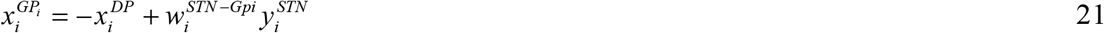

Since D1R is activated at increased dopamine levels, higher dopamine levels favor activating DP (constituted by the projections of D1R MSNs) over IP. This is consistent with the nature of switching facilitated by DA in the striatum (Chang et al., 2002; Frank and Claus, 2006; Lauwereyns et al., 2002; Tanaka et al., 2006). The relative weightage of the STN projections to GPi is represented by *w*^STN-GPi^, and is set to 1 for all the GPi neurons in the current study.

#### Choice Selection at Thalamus

GPi neurons project to thalamus through inhibitory connections. Hence the thalamic afferents can be simply expressed as a modified form of eqn. (21).

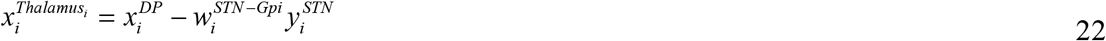

These afferents in eqn. (22) activate thalamic neurons as follows,

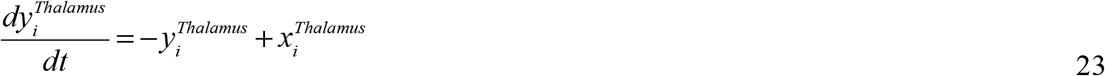

where 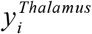 is the state of the *i*^th^ thalamic neuron. Choice selected is simply the ‘*i*’ (*i*=1,2,..,*n*) whose 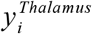 first crosses the threshold on integration. The threshold value used in the current simulation is 1.815.

### Reduced dynamical system model (Model C)

A simple dynamical systems approximation of the positive and negative attractor system described by models, A and B, can be given by:

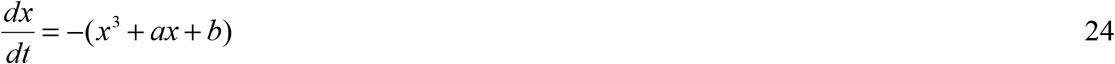

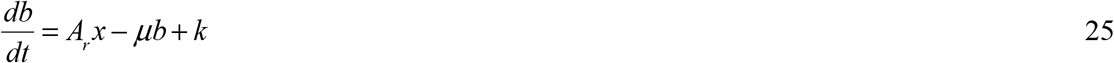

Eqn. 24, in isolation, shows bistability: for a large positive (negative) ‘*k*’, *x* stabilizes at a negative (positive) value. Note that ‘*b*’ tends to increase (decrease) for positive (negative) ‘*x*’ (eqn. 25).

The positive and negative wells generated by the x-cubic equation (eqn. 24) emulate the positive and negative utility states modeled by the critic component of previous models. The b-equation (eqn. 25) helps in setting up the stability of solutions, approximates the auxiliary dynamics (eqns. 18,19) of the previous models (A, B).

### Format of results

In all models A, B and C, we present the stability of solutions as a function of key parameters driving the models (*A*_r_ and *k*). Monostable and oscillatory solutions are presented for each model, where monostability is the presence of either positive or negative state attractor, and oscillatory is the presence of oscillations between positive and negative states.

For models A and B, the choice selections through time are smoothened by averaging with a moving window of size 50 (i.e., by averaging moving boxcars) to compute the percent of positive state as a choice at a given time. The trajectory of positive state selection percentages through time (read-out) is then fit with a polynomial of degree 50 to enable maximal good fit in MATLAB. The frequency of read-out fitting curve is found by first subtracting out their mean, then finding the index of absolute maximum in frequency (FFT) space. Multiplying with 365 divided by length of the read-out normalizes the index. The choice of numbers used for normalizing the index helps in presentation of results comparable with other models. Trajectories with frequencies greater than 1 in a period of 365 time units are taken to possess oscillations in read-out spaces. But, the oscillations between positive and negative state regimes are of key interest. To filter those trajectories, we compartmentalize the percent positive state selection greater than 50% to be in positive state regime, and that less than 50% to be in negative state regime. The read-out trajectories showing oscillations between positive and negative state regimes are then labeled to show bipolar oscillations; those that do not exhibit oscillations are labeled to show monostable solution. In the dynamical system model (C), the sign of ‘*x*’ variable determines the valence of states. Bipolar oscillations are defined by oscillations between solutions of *x* with opposite signs through time as computed in the read-outs. Monostability is interpreted when the choice read-out trajectories through time converges to a single state regime (positive or negative ‘*x*’). The stability of solutions in the reduced simple model (C) is computed analytically, by solving the cubic using Cardano’s method (Confalonieri, 2015), and mapping the resulting eigenvalues of their Jacobian (at equilibrium point) to the respective dynamical solutions.

## Results

### Model A

First, we describe softmax policy based phenomenological model A, set to exploitative mode with *β* =10 in eqn. (9), for selecting the highest utilities of positive and negative states. Dynamics in *α*, eqns. (7, 8), leads to an increase (decrease) in value of *α* when in positive state (negative state) due to the reward average magnitudes; this promotes the selection of negative state (positive state) as the utility of positive state (negative state) gradually reduces. The above mechanism causes oscillation between states, for certain values of reward sensitivity *A_r_*, and basal risk sensitivity or tonic serotonin level parameter *k*. The variation in periods of positive and negative cycles can also be controlled by other parameters such as *τ* _r_ and *τ* _a_ in the eqns. (7, 8); for a fixed value of *τ* _r_ = *τ* _a_ = 100, the results are as shown in figure 3. Figure 3 (first panel) portrays the stability of trajectory of read-outs (percent positive state selection through time) as a function of parameters, *A_r_* and *k*. Cases a (controls), b, and c of figure 3 show smoothened (window size of 20) trajectories of 13 different initial values of *α*.

**Figure 3-.**
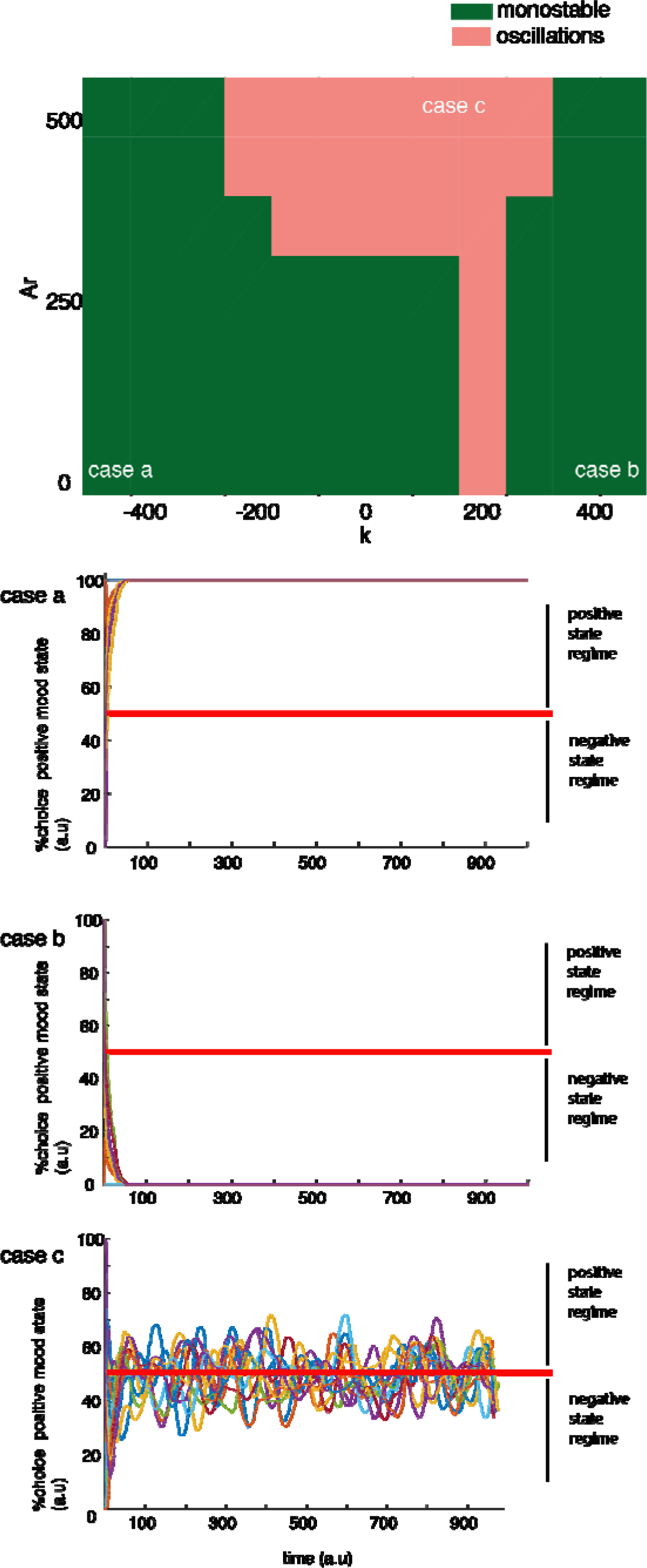
Softmax-based phenomenological model: The topmost panel shows the stability of solutions as a function of *A_r_* and *k*. Monostable solution could indicate either positive or negative state stability, and oscillations indicate swings between positive and negative states with varying time periods. Instances of each solution are provided as *cases a, b, c*: Solutions stabilizing at positive state regime as shown in case (a) for parameters *A_r_* = 0.001 and *k* = −500, and solutions stabilizing at negative state as in case (b) are shown for parameters *A_r_* = 0.001 and *k* = 500. Oscillations, case (c), are shown for parameters *A_r_* = 100, and *k* = −0.001.

### Model B

Next, we show oscillations between positive and negative states using a more realistic network model of BG working with the *α* dynamics as described in eqns. (18, 19). In this model, eqn. (11), value is represented by the activity of D1 MSN, and risk by D1-D2 co-expressing MSNs. The functional principles of computing the utility and selecting the maximum between utilities of positive and negative states remains as the same as the model A. The *α* changes as a function of reward averages, like that described in model A, to facilitate oscillations between states. Figure 4 (first panel) portrays the stability of trajectory of read-outs (percent positive state selection through time) as a function of parameters, *A_r_* and *k* Cases a, b, c of figure 4 show smoothened (window size of 20) trajectories of 13 different initial values of *α*.

**Figure 4-.**
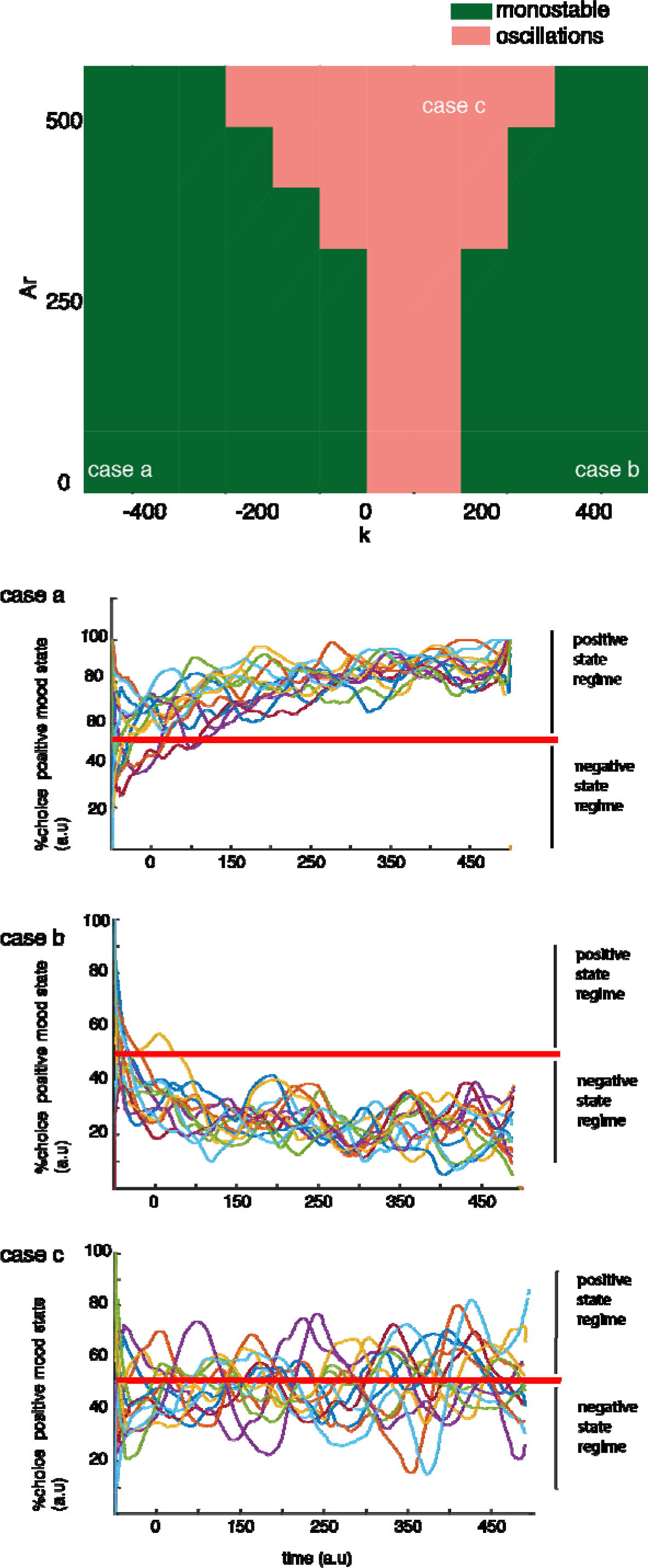
BG network model: The first panel shows the stability of solutions as a function of *A*_r_ and *k*. Monostable solution could indicate either positive or negative state stability, and oscillations indicate between among positive and negative states with varying time periods. Instances of each solution are provided as *cases a, b, and c*: Solutions stabilizing at positive state regime as shown in case (a) for parameters *A*_r_ = 0.001 and *k* = −500, and solutions stabilizing at negative state as in case (b) are shown for parameters *A*_r_ = 0.001 and *k* = 500. Oscillations, case (c), are shown for parameters *A*_r_ = 100, and *k* = −0.001.

### Model C

Furthermore, we capture the positive and negative regime attractors simulated using utility models A and B using a simple 2-dimensional model. This model uses a cubic ‘*x*’-variable equation to capture the positive (negative) attractors, while *x* is positive (negative), eqn. (24), respectively. The auxiliary dynamics (as simulated using α in models A and B) with key parameters *A*_r_ and *k* are captured in the equation simulating ‘*b*’-dynamics, eqn. (25). Analysis with the reduced model shows that the parameter representing reward sensitivity in the models A and B, i.e., *A*_r_, is equivalent to the coefficient of *b*-variable; and the basal risk sensitivity, i.e., *k* parameter, is equivalent to a constant that adjusts the height of the intersection point of *b* and *x* nullclines. Hence a negative (positive) *k* produces the intersection point at positive (negative) *x*, and thereby stabilizes the positive (negative) states. Furthermore, higher values of *A*_r_ facilitate limit cycle oscillations. Figure 5 (first panel) portrays the stability of trajectory of read-outs as a function of parameters, *A*_r_ and *k* The stability of solutions is computed using bifurcation analysis (as described in the methods). In the bistable case, the solution is dependent on the initial condition, is indicated by grey color in the result figure. Cases a, b, c of figure 5 show smoothened (window size of 20) trajectories of 13 different initial values of ‘*x*’ and ‘*b*’.

**Figure 5-.**
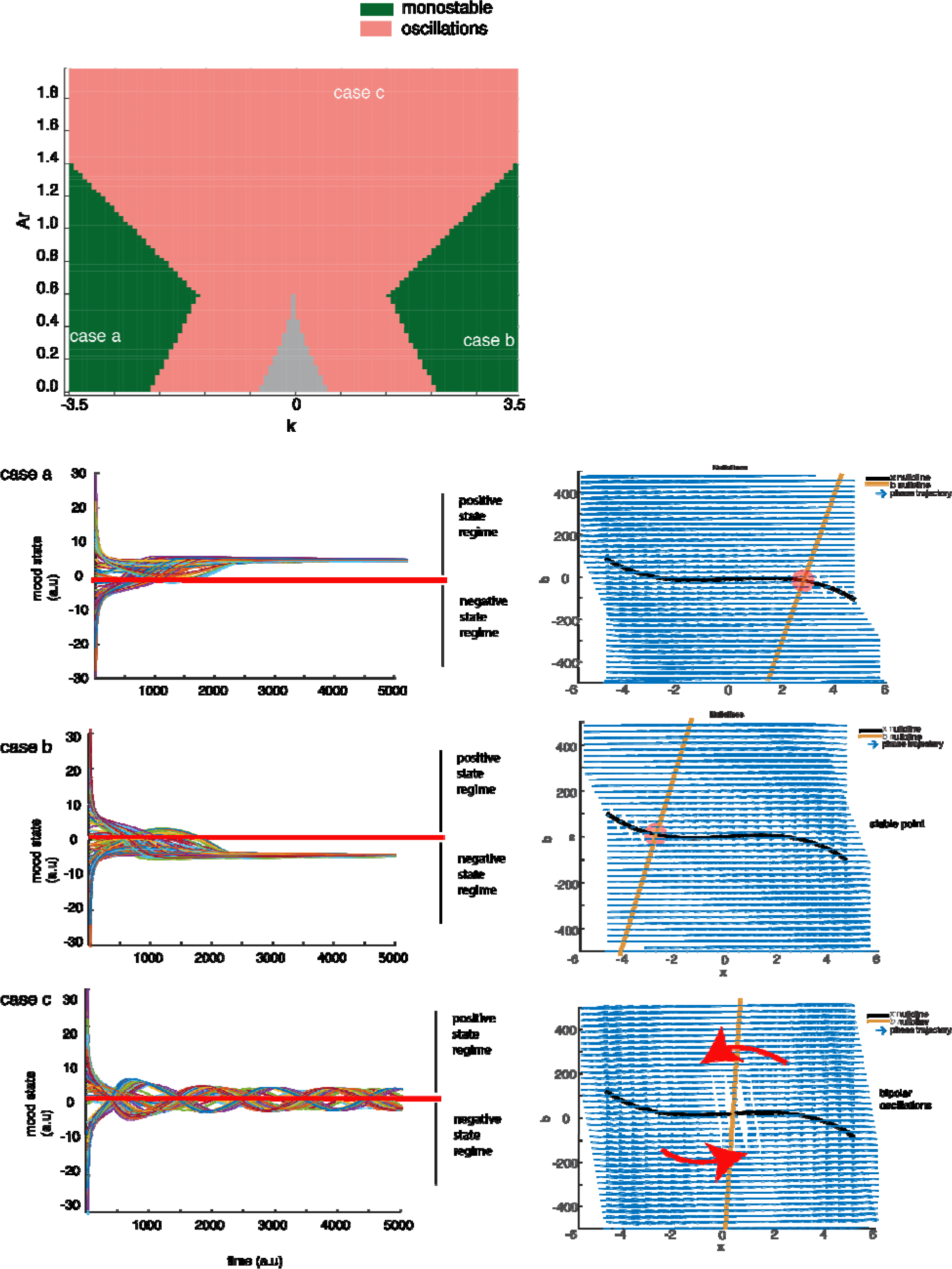
Reduced dynamical system model: The first panel show the stability of solutions as a function of Ar and k, with other parameters set to a = −6, and μ = 0.6. Monostable solution could indicate either positive or negative ‘*x*’ stability, and oscillations indicate swings between positive and negative states with varying time periods. Instances of each solution are provided as *cases a, b, and c*: Trajectories and phase planes for dynamics stabilizing at positive state regime as shown in case (a), at a negative state are shown in case (b), oscillations in case (c).

## Discussion

Bipolar disorder is characterized by mood swings, - oscillations between manic and depressive episodes, - with the episodes varying from hours to years (Hilty et al., 2006). The underlying factors of bipolar disorder are thought to involve a combination of genetic, biological, and environmental factors (Alloy and Abramson, 2010; Alloy et al., 2015; Harvey, 2008). We are interested in understanding the pathophysiology from the view of decision making dynamics. We explore the computational grounds facilitating occurrence of positive and negative stable mood states in an alternative manner. Particularly, we focus on a key factor contributing to the disorder namely dysfunction of the serotonergic and dopaminergic system in the reward circuitry mediated by cortico-basal ganglia network dynamics. To this end, our model finds that reduced risk sensitivity levels (*k*), and abnormally high reward sensitivity (*A*_r_) as key factors that contribute to alternating choices between manic and depressive episodes. A comparison with the reduced dynamical system show their correspondences to a simple system (model C), which can pave the way to understanding of the factors causing bipolar disorder in human patients.

Existing models use abstract limit cycle systems to show bipolar oscillations (Odgers et al., 2009), but they do not make a contact with the underlying neural substrates. We too begin with an abstract utility based softmax model of bipolar oscillations (model A), but expand it to a neural network model of BG (model B) that builds on our earlier modeling effort describing the roles of dopamine and serotonin neuromodulation in the decision making functions of the BG system (Balasubramani et al., 2014, 2015a). Under control conditions (case a of figures 3-5), the network selects the rewarding choice with a high probability. A two-variable reduced model C of the dynamics allowed exploration of the entire phase plane as a function of the two parameters of interest viz., reward sensitivity (*A*_r_), and basal risk sensitivity (*k*). A comparison between simple dynamical system and cortico-basal ganglia network substantiate two crucial factors contributing to bipolar-like oscillations of the model: *A*_r_ and *k* of the reduced model that correspond to the network model’s reward sensitivity (*A*_r_), and basal risk sensitivity (*k*), respectively.

There has been a lot of clinical and experimental evidence supporting 5-HT dysfunction and reward hypersensitivity in bipolar disorder (Hilty et al., 2006). Our modeling study suggests that reward hypersensitivity (Alloy et al., 2015) with medium levels of 5-HT as tonic/basal values or that induced by medication, can facilitate bipolar oscillations. Serotonin signaling has been linked to reward magnitudes and reward processes by various experiments (Kranz et al., 2010; Nakamura, 2013; Nakamura and Wong-Lin, 2014), and is influenced by habenular and PFC inputs (Challis et al., 2015), a fact that supports our model’s proposal to bidirectionally relate rewards to serotonin-mediated auxiliary dynamics (eqns. 18, 19). Moreover, our model links 5-HT levels to risk sensitivity, just as previous studies have suggested that risk-aversion and risk-seeking are altered in bipolar disorder (Chandler et al., 2009). A major pharmacological therapy for bipolar disorder is administration of lithium (Geddes and Miklowitz, 2013). There have been reports that lithium affects the sensitivity and function of serotonin receptors (Price et al., 1990; Wood, 1985). Hence controlling the sensitivity of 5-HT receptors has been shown to contribute to stabilize the moods and control the episode symptoms (Dremencov et al., 2005; Wood, 1985), just as predicted by our model. Along with 5-HT, DA control of the network has been proposed to relate to depression and manic disorders. There have been several proposals suggesting an increase of dopamine in manic state and a decrease during depression (Cookson, 1985; Cools et al., 2011; Huys et al., 2015), and therefore it is plausible that the main regulators of the network-DA and 5-HT - have been also involved in bipolar disorder manifestation (Berk et al., 2007; Geddes and Miklowitz, 2013; Mahmood and Silverstone, 2001; Silverstone, 1985).

Therefore, we model bipolar oscillations as a manifestation of impaired reward-based decision making framework, and describe mania and depression as distinct cognitive states with opposite reward outcomes. The reduced dynamical system model shows the similarity between analysis of our model to other catastrophe theory based models developed for disorders such as anorexia (Zeeman, 1976). Such an understanding supported by the proposed cortico-basal ganglia model may form a preliminary basis to pinpoint underlying neural dynamics for various pathological approach behaviors, and may assist in designing and interpreting the system level dynamics used for therapeutics (Alloy and Abramson, 2010) and cognitive behavioral therapy. Importantly, bipolar manic and depressive people have exaggerated cognitive scores of reward sensitivity, a feature observed in our model (Alloy et al., 2015). Similarly, the bipolar patients have impaired neuromodulatory control too (Berk et al., 2007; Geddes and Miklowitz, 2013; Mahmood and Silverstone, 2001; Silverstone, 1985). The scope of recurrence depends on the initial state of the system, to which therapeutics is provided to alter the internal dynamics. We propose that the systems level understanding of bipolar oscillation dynamics, as in this study, can contribute prominently towards understanding at multiple scales; it may be a better way to proceed with the problem of recurrences in a precise and personalized manner. Moreover other factors such as life style (working schedule) and circadian rhythm driven internal cycles might influence the onset and persistence of symptoms of bipolar disorder (Alloy et al., 2015).

